# Adult-born granule cells support pathological microcircuits in the chronically epileptic dentate gyrus

**DOI:** 10.1101/2020.05.01.072173

**Authors:** FT Sparks, Z Liao, W Li, I Soltesz, A Losonczy

## Abstract

Temporal lobe epilepsy (TLE) is characterized by recurrent seizures driven by synchronous neuronal activity. The dentate gyrus (DG) region of the hippocampal formation is highly reorganized in chronic TLE; in particular, pathological remodeling of the “dentate gate” is thought to open up pathological conduction pathways for synchronous discharges and seizures in the mesial temporal lobe. However, this pathophysiological framework lacks a mechanistic explanation of how macroscale synchronous dynamics emerge from alterations of the DG at the microcircuit level. In particular, the relative contribution of developmentally defined subpopulations of adult-born (abGCs) and mature (mGCs) granule cells to epileptiform network events remains unknown. To address this question, we optically recorded activity dynamics of identified populations of abGCs and mGCs during interictal epileptiform discharges (IEDs) in mice with chronic TLE. We find that disjoint subsets of IEDs differentially recruit abGC and mGC populations. We used these observations to develop a neural topic modeling framework, under which we find that the epileptic DG network organizes into disjoint, cell-type specific pathological ensembles, a subset of which are recruited by each IED. We found that statistics of this ensemble structure are highly conserved across animals, with abGCs disproportionately driving network activity in the epileptic DG during IEDs. Our results provide the first *in vivo* characterization of activity dynamics of identified GC subpopulations in the epileptic DG, the first microcircuit-level correlates of IEDs *in vivo*, and reveal a specific contribution of abGCs to interictal epileptic events.

**Highlights:**

- We relate electrographic signatures of epilepsy to microcircuit dynamics at single-cell resolution
- The chronically epileptic dentate gyrus granule cell network is organized in lineage-specific pathological ensembles
- A novel generative model framework for ensemble recruitment relates local field potential signatures to microcircuit activation during interictal epileptiform discharges
- Adult-born granule cell-dominated ensembles are disproportionately represented among the inferred ensembles
- The most active ensemble during an interictal epileptiform discharge can be decoded directly from the local field potential spectrum
- This Latent Ensemble Recruitment model of cell recruitment by interictal events is the first application of Bayesian topic modeling to *in vivo* two-photon calcium imaging data

## Introduction

Temporal lobe epilepsy (TLE) is a common neurological disorder characterized by recurrent focal seizures originating in the mesial temporal lobe, most commonly the hippocampus (HPC). While the circuit mechanisms of TLE are not well-understood, it is thought that the pathological features of TLE emerge from a complex series of microlevel reorganizational steps leading to persistent perturbation of the excitatory and inhibitory processes regulating entorhinal-HPC interactions (Muldoon et al., 2015, Feldt Muldoon et al., 2013, Bragin et al., 1999a, Uhlhaas and Singer, 2006). Hypersynchronous macrolevel network events associated with TLE, in turn, are thought to be primarily the result of such microlevel alterations to underlying network connectivity within the HPC formation (Netoff et al., 2004, Percha et al., 2005, Dyhrfjeld-Johnsen et al., 2007, Morgan and Soltesz, 2008, Farrell et al., 2019).

Macrolevel synchronous events in epilepsy can be identified in electroencephalography (EEG) or local field potential (LFP) recordings as large amplitude and long duration electrographic seizures (Gibbs et al., 1935, Penfield, 1939), high frequency events (Urrestarazu et al., 2007), and most commonly and frequently, interictal epileptiform discharges (IEDs) (Dichter, 1997, Staley et al., 2011, McCormick and Contreras, 2001). These EEG events are thought to arise from pathophysiological changes to excitatory and inhibitory microcircuits in the epileptic network (Dudek and Staley, 2007). IEDs are transient EEG events characterized by a short duration (<100 ms) and large amplitude, and include multiphasic discharges as well as single interictal spikes (Huneau et al., 2013). At the macrolevel, these events appear to be the result of widespread, recurrent, synchronous population firing, though recent investigations into population dynamics at the circuit level have revealed heterogeneity of individual neuron responses during IEDs (Feldt Muldoon et al., 2013, Keller et al., 2010). This suggests that relatively sparse sub-ensemble dynamics support IEDs, and that these dynamics may contribute to the diversity of epileptiform activity identified in EEG or LFP recordings. These observations have been corroborated in human epilepsy patients via multiunit activity recordings from depth electrodes (Schevon et al., 2019). In addition, observations which support this hypothesis have been made in animal models of TLE by imaging excitatory granule cells (GCs) of the dentate gyrus (DG) in an *in vitro* slice preparation (Feldt Muldoon et al., 2013), and imaging either excitatory or inhibitory cell types in CA1 *in vivo* (Muldoon et al., 2015). While both excitatory and inhibitory cell classes have been shown to be involved in IEDs, how the collective activity of heterogeneous principal cell types gives rise to local network dynamics during pathological activity in chronic epilepsy *in vivo*, especially in the upstream nodes whose outputs shape CA1 excitation-inhibition dynamics, remains incompletely understood.

As the entry node of the HPC, the DG plays a critical role in cognitive processing by regulating the propagation of HPC feedforward drive (Hainmueller and Bartos, 2020). At the microcircuit level, strong local inhibition and lack of recurrent excitation of GCs together result in the sparse GC population activity that is thought to support computational functions such as pattern separation and novelty detection (Chawla et al., 2005, Leutgeb et al., 2007). This intrinsic low excitability is also thought to enable the DG to restrict the relay of synchronous activity from the entorhinal cortex into the HPC, thereby regulating the propagation of excitatory activity. This property is called dentate gating (Heinemann et al., 1992, Lothman et al., 1991), and the breakdown of the DG gate is hypothesized to contribute to epileptogenesis, leading to seizure generalization as well as cognitive and memory deficits (Thind et al., 2008, Krook-Magnuson et al., 2015). In the adult DG, new GCs are continually generated and functionally integrated into DG circuitry (Toni et al., 2007, Toni et al., 2008, van Praag et al., 2002). This adult-born GC (abGC) subpopulation of the DG network is especially sensitive to seizure-induced reorganization (Althaus et al., 2019, Buckmaster and Schwartzkroin, 1994, Buckmaster and Dudek, 1999, Althaus et al., 2016, Danzer, 2018, Danzer, 2019), that may take the form of an altered level of neurogenesis (Varma et al., 2019, Parent et al., 1997, Cho et al., 2015), mossy fiber sprouting (Dudek and Sutula, 2007, Sutula et al., 1988, Tauck and Nadler, 1985, Scharfman et al., 2003, Althaus et al., 2016, Zhou et al., 2019), abnormal formation and persistence of basal dendrites on abGCs (Buckmaster and Schwartzkroin, 1994, Murphy and Danzer, 2011, Ribak et al., 2000, Buckmaster and Dudek, 1999) as well as ectopic dispersion and migration of abGCs into the hilus or CA3 (Scharfman et al., 2000, Houser et al., 1992, Althaus et al., 2019). A recent *in vivo* study of normal DG function demonstrated that abGCs are intrinsically more active and less stimulus-selective than their mature counterparts (mGCs) (Danielson et al., 2016a). Together, these findings strongly implicate abGCs in the development of epilepsy, and support a hypothesis in which the relative hyper-activity of abGCs is amplified through recurrent microcircuits in the epileptic DG, resulting in generalized synchronous seizure activity in TLE (Buckmaster and Schwartzkroin, 1994, Knight et al., 2012, Patel et al., 2004, Hester and Danzer, 2014, Danzer, 2018, Danzer, 2019). However, at present there are no *in vivo* data allowing direct comparisons of the activity patterns of major GC subpopulations and their relative contribution to functional network structure during macrolevel epileptic events.

In the present study, we genetically label populations of mGCs and abGCs in the HPC before inducing the intra-HPC kainic acid (KA) model of chronic TLE, and then dissect the activity dynamics of these neural populations using a combination of *in vivo* two-photon calcium imaging and LFP recording. To uncover ensemble structure underlying high-density calcium recordings of the chronically epileptic DG network, we introduce a novel generative model framework for ensemble recruitment that captures existing knowledge and biological intuitions about the mechanisms of IEDs. Performing statistical inference on this class of model is difficult in general, but we show that our biologically motivated model can be reduced to Latent Dirichlet Allocation (LDA), a well-known topic model with many tractable inference algorithms (Blei et al., 2003). We use this reduction, in combination with tools from the LDA literature, to infer the recruitment of micro-ensembles by network-level events. Consistent with previous functional imaging in acute *in vitro* DG slices from chronically epileptic animals (Feldt Muldoon et al., 2013), we found structured ensemble dynamics within the GC population during IEDs *in vivo*. In addition, we also identified cell-type specific ensemble structure nested within the abGC and mGC populations, with distinct contributions to network dynamics in the interictal period. Finally, we found that abGCs participate in network activity during IEDs to a disproportionately higher degree than mGCs. These observations suggest that the chronically epileptic DG network exhibits robust underlying functional organization in temporal correlation and GC lineage. Because abGCs are strongly reorganized in TLE compared to their mature counterparts (Althaus et al., 2019, Buckmaster and Schwartzkroin, 1994, Buckmaster and Dudek, 1999, Althaus et al., 2016), our results suggest that this reorganization produces functionally distinct ensembles that have different pathophysiological roles identifiable *in vivo*.

## Results

### Two-photon calcium imaging of the epileptic DG *in vivo*

Dissecting the functional and anatomical microcircuit structure of the DG *in vivo* requires techniques that allow simultaneous recording of neural activity from a wide section of the network at single-cell resolution, while also permitting molecular identification of the recorded cells. This was achieved by performing two-photon calcium imaging of the DG in head-fixed mice on a circular treadmill (Danielson et al., 2016a). Combining these techniques with a cell-type specific *Cre* driver line allowed us to indelibly label and image abGCs and mGCs simultaneously in a single recording (Danielson et al., 2016a) (**Figure 1A**,**B**), which is not feasible with electrophysiological recordings or in freely moving animals. To indelibly label abGCs, NestinCreER^T2^ mice were crossed with a conditional reporter line (Ai9) and pulsed with tamoxifen (TMX) to express the fluorescent red reporter (tdTomato) in young abGCs (abGCs were 5 weeks old at the start of imaging, **Figure 1C**, see **Methods**). Mice were then stereotactically injected in the dorsal DG with a recombinant adeno-associated virus (rAAV) to express GCaMP6f in all GCs. KA was injected unilaterally into the ventral HPC ipsilateral to the viral injection site to induce *status epilepticus*, simulating the initial insult that leads to epileptogenesis in humans (Levesque and Avoli, 2013, Zeidler et al., 2018). The KA injection site was chosen to minimize associated gliosis and cell loss within the imaging field of view, thereby permitting imaging of the DG within the vicinity of the main insult to the network (Zeidler et al., 2018). Three days later, a chronic imaging window was implanted over the dorsal DG providing optical access necessary for cell-type-specific imaging of the GC layer in the dorsal blade of the DG. An ipsilateral single-channel LFP electrode was also implanted in the HPC ipsilateral to KA injection targeting the DG, and an electromyographic electrode implanted in the cervical trapezius neck muscle (**Figure 1B**,**C**). After recovering from the implant surgery, mice were continuously video-EEG monitored, and TLE onset was determined by the occurrence of the first detected generalized motor seizure event, which occurred on average 10 days after KA injection (242.9±20.4 hours, mean ±s.d.) in n=5 mice.

**Figure 1.**
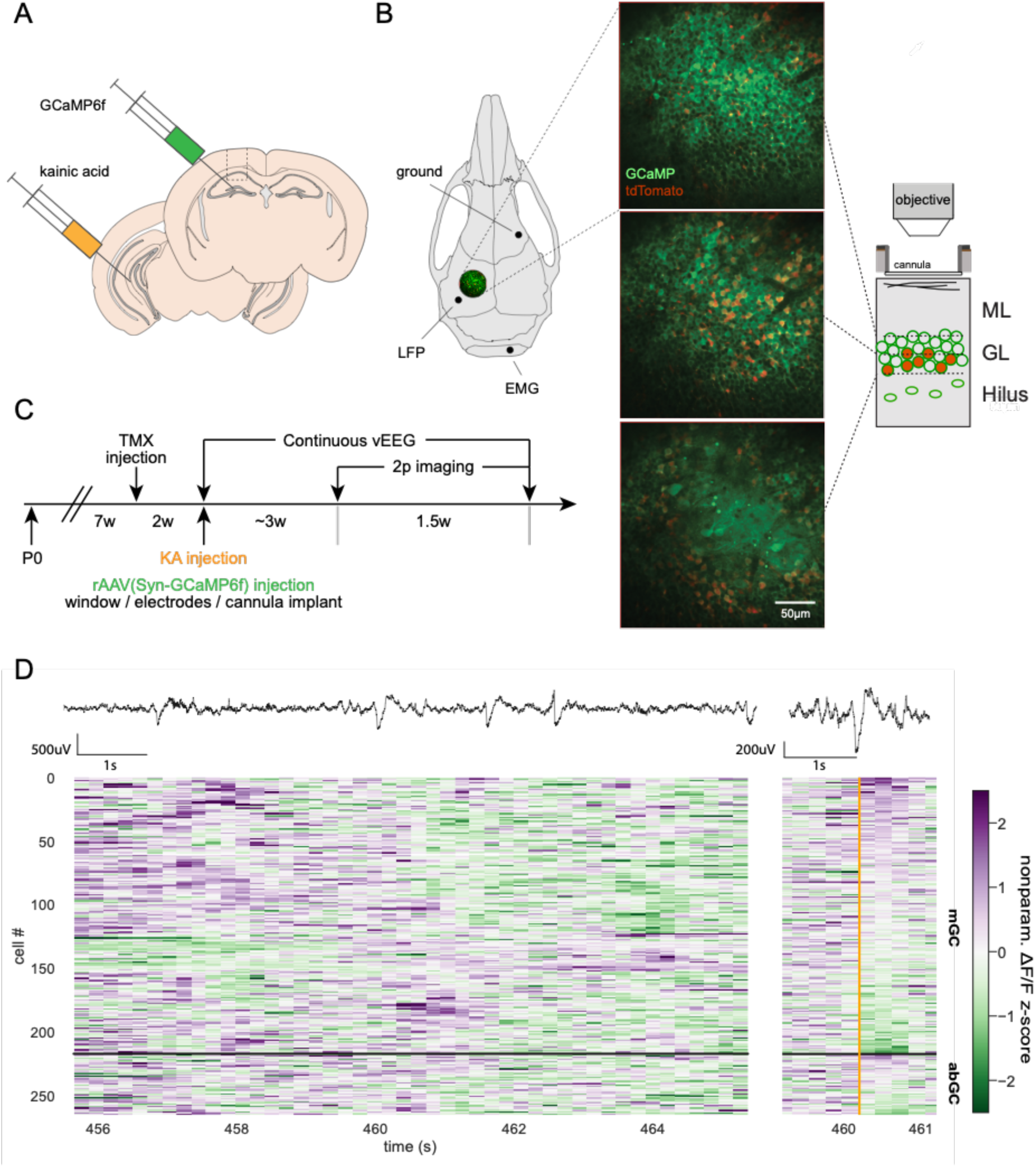
Cell-type specific functional imaging in the dentate gyrus. **A.** Experimental schematic. GCaMP6f is virally expressed in all DG neurons at the dorsal HPC site of injection, and kainic acid is injected into the ipsilateral ventral HPC to induce the model of epilepsy. **B.** Left: Schematic of mouse skull showing locations of imaging window, ipsilateral local field potential (LFP) depth electrode, electromyography (EMG), and ground electrode placements. Center: Two-photon line-scanning microscopy allows for the recording of large populations of GCs in surgically exposed dorsal DG. Representative example in vivo two-photon two-color microscopy time average images from near simultaneous multi-plane imaging in the mouse DG. The top two planes are focused in the dorsal GC layer while the bottom plane is located just above the polymorphic hilar region. Green represents GCaMP fluorescence in all cell types, while red represents tdTomato expression limited to abGC in a Cre-dependent manner. Right: Schematic showing near-simultaneous multiplane imaging throughout the DG granule cell layer. ML: molecular layer, GL: granule cell layer. **C.** Experimental timeline for the labelling of abGCs, induction of the kainate model, and two-photon imaging of the epileptic DG network. Adult-born GCs were indelibly labelled with tdTomato following injections of tamoxifen (TMX) that drove expression of Cre in Nestin+ cells. Two weeks later, kainic acid (KA) was injected into the ventral HPC and GCaMP6f was injected into the dorsal DG and chronic imaging window cannula was implanted above the dorsal DG. Following injection of KA, mice were monitored for ictal activity using continuous video-EEG. **D.** Top: Filtered LFP showing interictal epileptiform discharges in an IHK mouse 31 days after KA injection. Due to variability in IED amplitude, only a subset of IEDs are visible at this scale. Left: Example abGC and mGC whole population activity during IEDs across a 10 second time window. Activity for each neuron is shown as a positive (purple) or negative (green) nonparametric *z*-scored ΔF/F, clipped to ±2.5z for visualization. Right: LFP trace and population activity from the same abGCs and mGCs centered on an example IED (orange) across a 2 second time window and sorted based on maximum activity.

Given that the laminar organization of GCs in the dorsal DG is parallel to the optical field of view (Amaral et al., 2007), we can capture a large number of GCs within the GC layer and the sub-granular zone (**Figure 1B**). To further increase simultaneous imaging of multiple cell-type specific populations across these layers, we coupled our image acquisition control to a piezoelectric crystal for fast axial focusing and near simultaneous multiplane imaging (Danielson et al., 2016b) (**Figure 1B**). With multiplane imaging, we were able to simultaneously capture on the order of 433±261 mGC and 56±31 abGC (mean ± s.d.) active regions of interest within a single subject (example LFP recording, population *ΔF/F* to series of IEDs and example traces triggered by one IED shown in **Figure 1D**, note that only a fraction of the GCs participate in any given IIS; see below). In total, across 5 mice, 2164 mGCs (tdTomato-negative) and 278 abGCs (tdTomato-positive) (**Table S1**) were identified as active GC regions of interest during three consecutive 10 min long imaging sessions for each mouse (see **Methods**).

### Time domain, frequency domain, and mean calcium response of IEDs

The low probability of capturing a spontaneous seizure event during a two-photon imaging session poses a challenge to the optical dissection of epileptic networks *in vivo*. However, IEDs provide a series of functional “snapshots” of discrete components underlying the ictogenic circuits, during which only a subset of the network is activated (Feldt Muldoon et al., 2013, Muldoon et al., 2015). Therefore, we sought to analyze network activations during IEDs with the aim of first understanding these “snapshots” individually, and ultimately assembling them into a complete tapestry of the network. To this end, we segmented the LFP into time bins matching the imaging frames, and classified the imaging frames during which IEDs occurred. Traditionally, IEDs are classified by hand, based on morphological properties such as event width, deflection amplitude, and deflection shape; however, annotating these features for each event manually is labor-intensive. We developed a graphical interface for semi-supervised IED classification and detection to generalize these principles (**Figure 2A**). Because all information about the width, amplitude, and morphology of each event is contained in its frequency-domain representation, we trained a classifier on the power spectrum of a small set of hand-identified IEDs and allowed it to generalize and update its predictions in an interactive manner; final classification results were verified for correctness by the experimenters (see **Methods**). Other types of interictal events, such as high-frequency oscillations were not included by the classifier and were thus excluded from further analysis (**Figure 2A**). Consistent with previous work (Staley and Dudek, 2006), we found morphological heterogeneity within the classified IEDs. Despite this heterogeneity, frames containing IEDs exhibit a stereotyped LFP power spectrum after applying a 1/f correction, supporting the reliability of this classification method (**Figure 2B**). We find an average IED rate of 1.13±0.53 IED/s, with an average of 680±319 IEDs per 10-minute recording session (mean ±s.d.) (**Figure 2D**). We then sought to quantify the modulation of single cells by IEDs. It has been hypothesized that IEDs are primarily driven by abGC activity (Iyengar et al., 2015), and indeed we found that abGCs are significantly more responsive to IEDs on average compared to mGCs (p<1×10^−4^, Mann-Whitney *U*-test), though both are very close to 0 (**Figure S1A**), without significant differences in the responses of the two populations to onset (p=0.39) or offset (p=0.08) of locomotion (**Figure S1A**). Nevertheless, we found that the mean IED-triggered PSTH was flat for both abGCs and mGCs (**Figure 2C**). This observation is important as it establishes that at least one of two conditions must hold: either (a) most IEDs only recruit a very sparse subset of cells (in which case the “average” cell’s response to any particular IED would be small), or (b) most cells only respond to a very sparse subset of IEDs (in which case any particular cell’s response to the “average” IED would be small). In the remainder of the paper, we show that not only are both (a) and (b) true, but we can also identify subsets of cells which are consistently recruited together, thereby relating macrolevel LFP activity to the activity of individual neurons in a local network.

**Figure 2.**
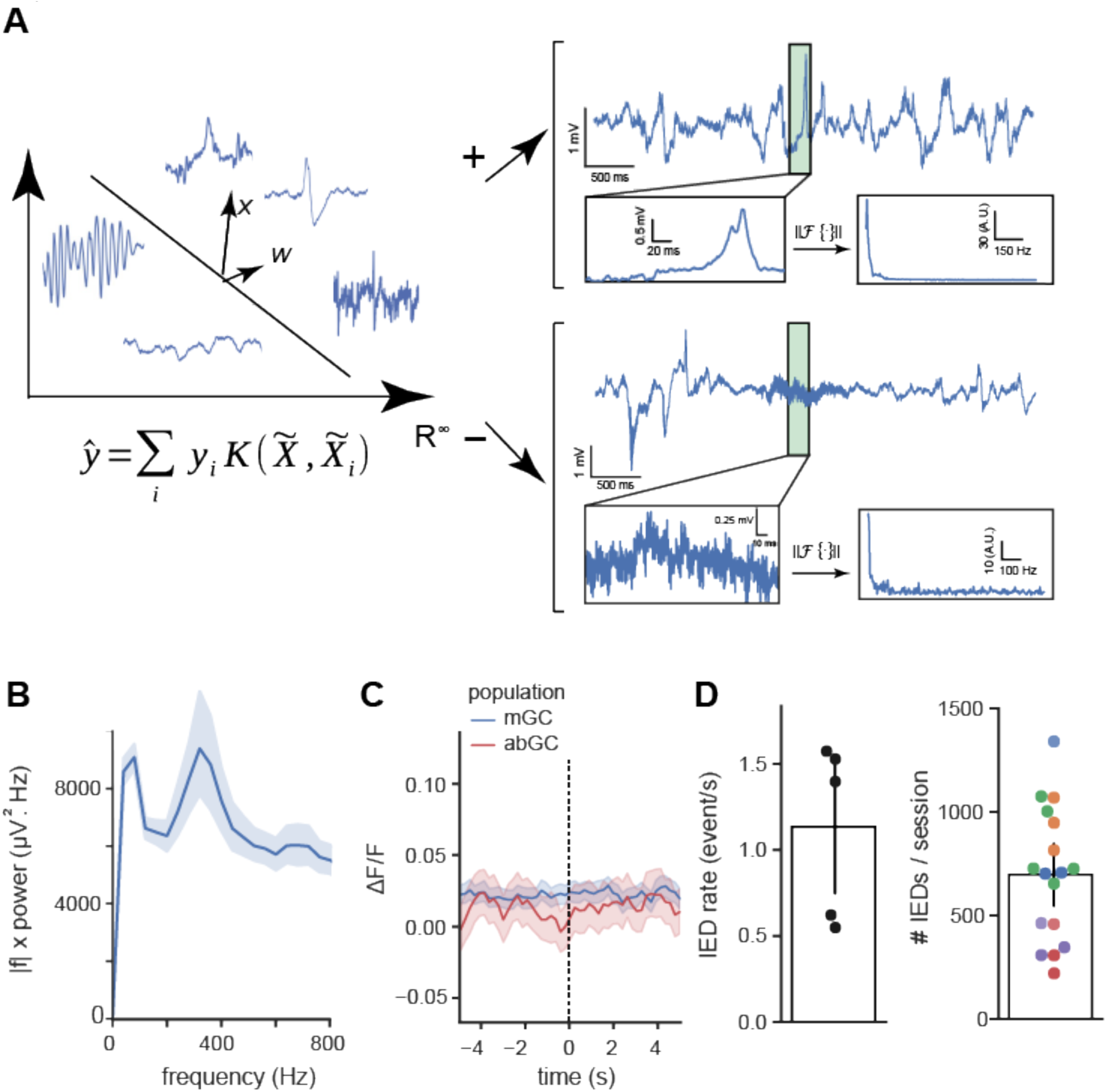
Interictal spike identification and basic characterization. **A.** Classifier schematic. Left: An Online Kernelized Perceptron was trained on the LFP magnitude spectrum where the window size was locked to each imaging frame. This algorithm was chosen because it permits efficient online training with immediate feedback after each labeled example and highly accurate phase-agnostic classification with a small number of samples. Right: Examples shown of classified interictal spike (above) and non-spike (below) highlighted in green, with zoomed-in view (below left) and magnitude spectrum (below right). **B.** Average IED magnitude spectrum. Average magnitude spectrum of n = 459 auto-detected events, with 1/f power correction applied. A peak frequency occurs in the 250-400 Hz range. **C.** Spike-triggered PSTHs showing averaged response to 4150 spikes by cell type, from 49 abGCs (red), 129 mGCs (blue) from one animal. The vast majority of GCs are *not* strongly modulated by any individual spike on average. **D.** Summary of detected IEDs rate across mice (n= 5 mice). Left: Average IED rate, in events per second (mean 1.13 IED/s, std 0.53 IED/s). Each point corresponds to a mouse (average of 3-5 sessions per mouse). Right: Total IED count per 10-minute recording session, mean count = 680 IEDs/session, std = 319 IEDs/session (each point corresponds to a single session, each color corresponds to a different mouse).

### Different IEDs recruit different populations of GCs

IEDs represent heterogeneous, global events in the epileptic brain (Bragin et al., 2000, Staley and Dudek, 2006). Given that we can only observe IEDs in a local network, treating them in aggregate as if they are homogeneous obscures the microlevel population activity patterns underlying each individual IED. Therefore, by training a logistic regression linear classifier to decode cell identity based solely on the cells’ IED responses, we assessed whether calcium responses of abGCs and mGCs differed on an IED-by-IED basis (**Figure 3A**). This classification procedure is equivalent to projecting the original *M*-dimensional responsiveness vector for each cell (where *M* is the number of electrographically identified IEDs) down to a 1D line representing a scalar “mGC-ness” score. The histogram of scores is bimodal, i.e., mGCs (blue) are mapped to positive scores and abGCs (red) are mapped to negative scores (**Figure 3B**). To perform the projection, a real-valued “weight” (A.U.) shared by all cells is calculated for each IED, which can be interpreted as the degree to which positive modulation by that IED is associated with mGC identity (positive weights, which we call a “pro-mGC” IED) *versus* abGCs (negative weights, “anti-mGC” IED). Under logistic regression, the two classes are assumed to be contrastive; that is a “pro-mGC” IED is associated with both positive modulation of mGCs or negative modulation of abGCs, and likewise an “anti-mGC” IED is associated with negative modulation of mGCs or positive modulation of abGCs. Sorting the weights (thresholded to a cutoff of ±0.1) by IED time (**Figure 3B**, middle) shows that most events are associated with some pro-mGC or anti-mGC bias, though this analysis does not permit us to say whether the activations are structured, as the sequence of mGC- and abGC-predictive events appears random when treating the two populations as internally homogeneous (**Figure 3B** middle and bottom). Thus, we conclude that each individual IED carries a small amount of information about cell identity, but the recruitment profile of each event is highly stochastic as any particular cell may be positively modulated, negatively modulated, or unresponsive to any particular IED regardless of the cell’s identity. However, the complete profile of a cell’s response to all IEDs is sufficient to predict cell identity (i.e., abGC or mGC) with high accuracy (**Figure 3B**).

**Figure 3.**
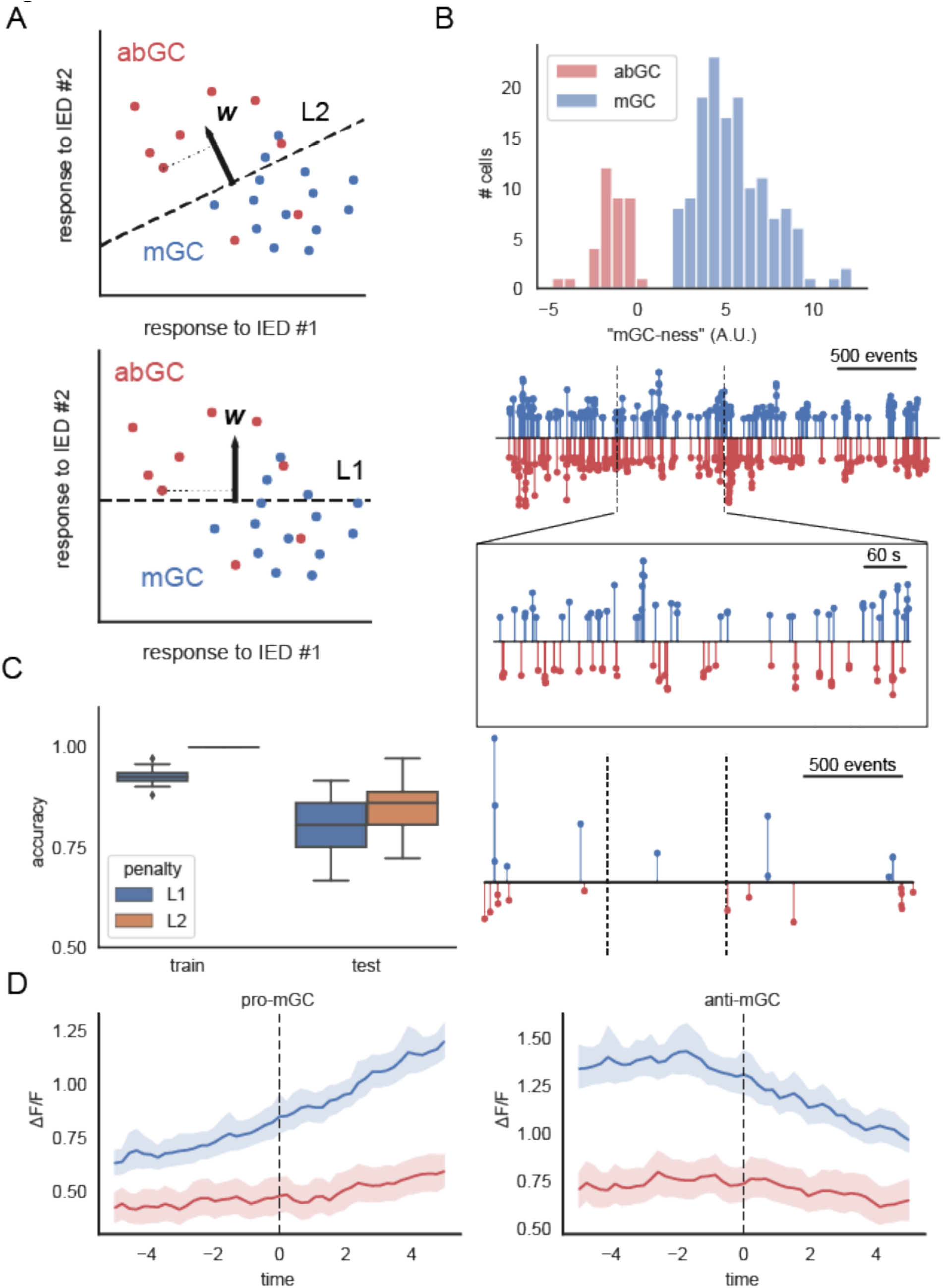
Linear classifier shows differential participation of abGC and mGC populations during interictal spikes. Considering each interictal event individually reveals that some spikes preferentially recruit abGC (red) or mGC (blue) neurons, using the convention “preferred” = positively modulated. **A.** Schematic of L1 and L2 embedding/classification procedure. Logistic regression projects the “response fingerprint” of every cell down to a maximally separating line in spike-response space (***w***). L1 regularization introduces sparsity to ***w***, effectively choosing the fewest events necessary to perform accurate classification. **B.** The 1D projection can be interpreted as an “mGC-ness” score (top histogram). The embedding weights (stem plot, A.U.), plotted in order by event number over a 30-minute recording session, quantify the degree of positive or negative mGC modulation by each event. Expansion: IEDs in the middle 10 minutes of the session, plotted against real time. **C.** Logistic regression on IED responsiveness with L1 (blue) and L2 (orange) regularization predict cell identity significantly better than chance. Logistic regression with L1 (“lasso”) regularization discovers a sparse subset of interictal spikes which is “most informative” about cell identity. Left: 100-fold cross-validation on an 80-20 train-test split shows 85±5% classification accuracy on the test set under standard L2 regularization and 81±5% classification accuracy under L1 regularization, implying that a small subsets of IEDs are highly associated with either mGC or abGC activation, such that cell activity within these events alone is sufficient to accurately predict cell identity. Right: Similar to panel *B*, weights associated with “most informative” events (A.U.) identified from lasso classifier plotted vs event number over 3 back-to-back recording sessions (vertical dashed lines). Events which are predictive of mGC identity (i.e., events where mGCs tend to exhibit positive responsiveness, or abGCs tend to exhibit negative responsiveness) termed ‘pro-mGC’ events, are colored in blue and point upwards, while events where mGCs tend to exhibit negative responsiveness or abGCs tend to exhibit positive responsiveness are termed ‘anti-mGC’ events, and are colored in red and point downwards. **D.** PSTHs triggered on pro-mGC (left) and anti-mGC (right) IEDs for an example mouse. Pro-mGC IEDs positively modulate mGCs, while anti-mGC events negatively modulate mGCs (right, in blue). In contrast, abGCs do not appear to be significantly modulated in either direction by either event type, suggesting that each IED may only recruit one population or another (or neither).

Based on the cells’ IED responses, the classifier achieved 85±5% accuracy on a test set, balanced to remove the effect of population size, cross-validated over 100 random 80-20 train-test splits (**Figure 3C**, left). Subsequently, we introduced a sparsifying (L1), regularization penalty in order to concurrently identify the subset of IEDs which are “most informative” about cell identity, in the sense that knowing a cell’s response to those events is almost as predictive of identity as knowing the cell’s complete response profile (**Figure 3C** right; see **Methods**). Analyzing the informative events identified by this regularized classifier will reveal how cell identity influences participation in IEDs. For instance, the regularized classifier could find that participation in a particular sequence of mixed events is predictive of abGC identity, which would indicate that abGCs and mGCs are internally coherent and respond to IEDs through population-specific patterns. A second possibility is that the regularized classifier may find individual “pure” or near-pure mGC-associated IEDs, which would indicate that some IEDs are attributable almost exclusively to one population. Finally, a third possibility is that the regularization penalty results in a much worse test accuracy, which would indicate that the entire profile of a cell’s response to IEDs is necessary to classify the cells accurately.

The L1 regularization penalty resulted in a slight reduction in test set accuracy (81±5% accuracy using the same procedure), but identified IEDs with highly population-specific recruitment profiles and disregarded the majority which exhibited mixed activation, suggesting the second hypothesis above holds true (**Figure 3C**; see **Methods**). Single cell responses to the “most informative” IEDs identified by this procedure showed high within-population heterogeneity but striking between-population differences (**Figure S1B**): in contrast to the flat undifferentiated IED-triggered response (**Figure 2C**), we found subsets of IEDs that significantly modulate the mGC population (both positively and negatively), while the abGC population is not significantly modulated in either direction by either event type (**Figure 3C, Figure S1C**). Thus, we conclude that most IEDs differentially recruit abGCs compared to mGCs, but the recruitment of any particular cell by any IED appears random. Reliance on only one population is allowable under the assumptions of logistic regression, as an algorithm that classifies from the mGC population with high sensitivity and specificity can achieve high accuracy overall by simply classifying “non-mGC” examples as abGCs. This limitation highlights the need for a model which can more expressively capture the activation of different populations, and perhaps functionally-defined subpopulations, by different subsets of IEDs.

### A novel generative model of latent ensemble recruitment uncovers within-population ensemble dynamics

The linear decoding analyses above implies heterogeneous, cell type-related dynamics among IEDs. However, we have so far assumed that abGCs and mGCs form internally homogeneous populations, while there is a growing body of evidence that this is not the case (Erwin et al., 2019). A limitation of logistic regression in this setting is that a mix of abGCs and mGCs respond to most IEDs, but the mix itself may be structured. Even among the “most informative” pro-mGC and anti-mGC IEDs identified by the sparse decoder (**Figure 3C**), there is clear heterogeneity of the responses between individual cells within a population, even within a single mouse (**Figure S1B**). Furthermore, the linear decoding analysis classified abGCs as those cells that did not significantly respond to pro-mGC or anti-mGC IEDs, whereas we might also like to identify IEDs that independently significantly modulate the abGC population. We sought to construct a generative model that relates the population activity in the imaged local network to macrolevel IEDs via the hidden functional ensemble structure within the network, which can be compared *post hoc* with ground truth GC identities. This model should be able to account for the ensemble structure and activation underlying the data with the following properties:

1. Identifies functional ensembles that can be compared with cellular identities
2. Infers underlying ensemble structure incrementally revealed by successive IEDs, even if an entire ensemble is never observed to be active all at once
3. Sliding scale of activation: Ensembles may be activated to differing degrees by different IEDs
4. Mixed membership: Cells may be associated with more than one ensemble, and may be associated to differing degrees to ensembles
5. Parameter inference (ensemble composition, activation in each IED) from data is computationally tractable

We developed a novel generative model motivated by Bayesian topic modeling (Blei et al., 2003, Hathway and Goodman, 2019), that we call Latent Ensemble Recruitment (LER), and that has all of these properties. This LER model (**Figure 4A, Figure S2A**) assumes that the network consists of *K* fixed, unobserved ensembles of cells. Each IED recruits a sparse subset of these ensembles, and each ensemble may activate a few or many of its cells, depending on the degree to which it was excited by that IED (see **Methods** for full description of the generative process). Various strategies are possible for performing inference on this model, though the number of latent variables we must learn compared to the number of observables presents a challenge. One convenient inference strategy is to perform *ad hoc* inference on *z* using bootstrapping to binarize the dataset (*z’*), in order to solve the resulting inference using a standard variational Bayes solver (Pedregosa et al., 2011). This *ad hoc* procedure is equivalent to reducing our model to Latent Dirichlet Allocation, a closely related model for which the inference problem has been well-studied, using the topic modeling analogies: ensemble ∼ “topic”, spike ∼ “document”, cell ∼ “word” (Blei et al., 2003) (**Figure S2A**). The binarized data *z’* (**Figure 4B**) contains some qualitative ensemble structure and temporally distinct activations between abGCs and mGCs. The ensemble activation matrix θ (**Figure 4C**) shows temporally coherent activations (i.e., an ensemble activated by event *i* is likely to also be activated by events *i* − *1* and *i* + *1*), despite the exchangeability of IEDs (i.e., the model receives no information about the order or relative timing of events). This temporal coherence may be due to temporary increases in excitability localized to the microcircuits underlying each IED, such that subsequent IEDs are more likely to recruit the previously excited ensembles. **Figure 4D** shows examples of learned ensembles σ: we say a cell “participates” in an ensemble *k* if the weight of that cell in s k is > 3/N (i.e., greater than 3x uniform prior level). Even though the model receives no information about cell lineage, from these inferred results we can clearly distinguish mGC-predominant, abGC-predominant, and mixed ensembles (**Figure 4D**). Consistent with results previously reported *in vitro* (Feldt Muldoon et al., 2013), we also find low overlap (≤ 3 cells) between any pair of ensembles, as measured by the Pearson correlation between ensemble vectors (**Figure 4E**, right). Because of this low overlap, we can interpret the LER results as a sort on the cell-cell correlation matrix. Sorting the correlation matrix accordingly, we find that the cells exhibit high correlations within ensembles and lower correlations outside their ensemble as expected (**Figure 4F**), providing independent confirmation that the ensembles discovered by the model are reasonable. Finally, we sought to determine whether the most active ensemble in each IED could be decoded from the power spectrum of the IED itself. For small ensemble number (K), we can decode the most active ensemble in a held-out test set significantly better than chance using a random forest classifier with 100 estimators trained on the LFP spectrum collected within an imaging frame (**Figure 4H**). We conclude that IEDs are driven by multiple ensembles with temporal and lineage-dependent structure, and conversely, different ensembles are associated with distinct electrographic signatures on LFP.

**Figure 4.**
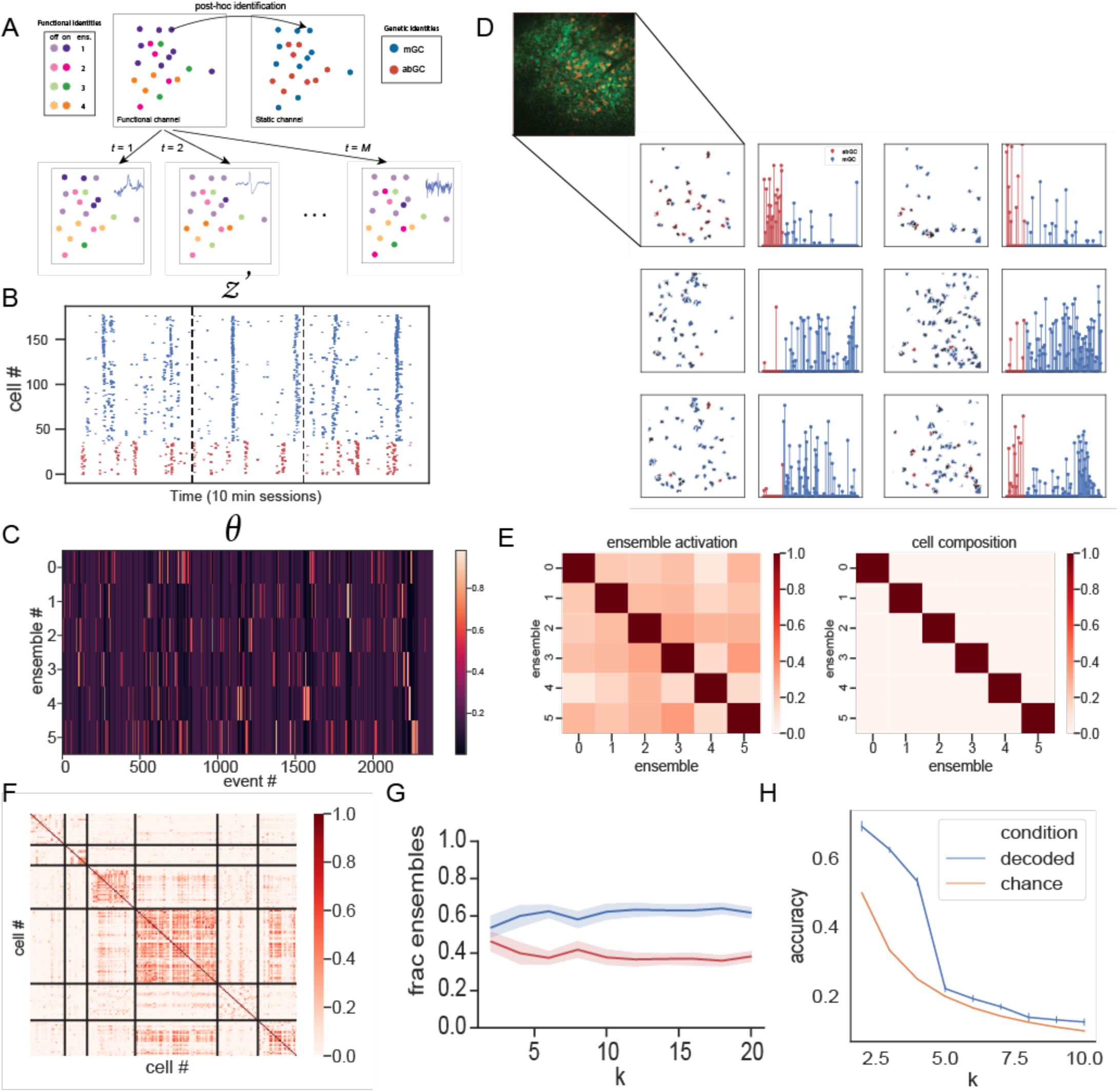
A generative model of spike ensemble recruitment. Latent Ensemble Recruitment (LER) model uncovers latent ensemble structure in the data. **A.** Schematic of LER generative model. Colors represent latent (hidden) ensembles in the data. Cells (colored circles) are colored by the ensemble they are most associated with. Cells in the same ensemble tend to be recruited together by IEDs (bottom), but recruitment is in general sparse and random. The hidden ensemble structure is revealed by many observations. The ensemble identities can be compared to known genetic identities *post hoc*. **B.** Raster plot of bootstrapped “activation” variable z 0, 10,000 bootstraps with α = 0.05, sorted by identity (red: abGCs, blue: mGCs) in synchronous network events excluding seizures in three consecutive 10 minute recording sessions, plotted against real time (n = 178 cells, n = 129 mGCs, n = 49 abGCs). **C.** Ensemble activation matrix, plotted against event number, ordered in time (K = 6 ensembles, M = 2473 events). The model is agnostic to the order in which events are presented to it, but nonetheless discovers temporally structured activations, e.g., ensemble 4 is repeatedly activated for a string of consecutive IEDs after event 1500. This suggests that while recruitment of individual cells may be random, the recruitment of ensembles exhibits statistical regularities which correspond to biological intuitions about the underlying circuits, e.g., a microcircuit may be excited and remain in a hyperexcited state for several seconds such that it is more likely to be recruited by global activity. See *Figure S2* for cross-validation across different mice and model fits. **D.** The six ensembles identified from data in *C*, with ensemble activations shown in *D*, with spatial distribution (left; overlay of 4 imaging planes), colored by weights (A.U., right). Each pair of plots represents two views of each ensemble – on the left, the spatial footprints of cells associated with that ensemble in the imaging field of view, (red = abGC, blue = mGC), and color intensity corresponding to the strength of their association with that ensemble (A.U.). On the right, each lollipop represents a cell, with the cells grouped by molecular identity and height (A.U.) representing the strength of each cell’s association with an ensemble. Clustering by molecular identity is readily apparent, with abGCs dominating the activity of ensemble 0, mGCs of ensembles 1, 2, and 4, and ensembles 3 and 5 mixed. See *4G and Figure S2* for cross-validation across different mice and model fits. **E.** Characterization of learned ensembles and activity patterns. Left: Ensemble activity patterns are weakly correlated, as measured by Kendall’s tau. In this context, Kendall’s tau can be interpreted as “bubblesort distance,” i.e., the amount of temporal distortion required to align two activity patterns. Our model is able to accommodate the joint activation of multiple ensembles in a single event; this observation suggests that inter-ensemble correlation is present in our data, but only weakly. Right: Ensembles are overwhelmingly mutually exclusive; although our model is able to accommodate mixed membership (i.e., cells which belong to multiple ensembles), the learned ensembles are close to orthogonal (Pearson’s correlation, p > 0.05), suggesting the epileptic network organizes into *disjoint* pathologic ensembles. **F.** LER learned ensembles as a sort on the cell-cell correlation matrix. The previous observation that ensembles are disjoint means that an unambiguous membership function exists from cells to ensembles, i.e., a cell can be said to “belong” to an ensemble without ambiguity. We sort the cells by ensemble (defined as the ensemble in which each cell achieves its highest weight), which reveals previously unseen structure in the Kendall tau correlation matrix. This shows that cells within a single ensemble have highly correlated activity patterns, while cells in different ensembles tend to have low correlation. **G.** Fraction of abGC vs mGC-dominated ensembles, plotted as a function of K. The ratio of abGC-dominated ensembles to mGC-dominated ensembles remains fixed at 4:6 across mice, suggesting that abGC-dominated ensembles disproportionately drive network activity. **H.** Decoding the most active ensemble from the LFP power spectrum of the imaging frame during which the IED occurred. For small values of the ensemble hyperparameter, we can decode the most active ensemble in a held-out test set significantly better than chance using a random forest classifier with 100 decision tree estimators trained on the LFP power spectrum, validated across 10 random 80-20 train-test splits. As *K* is increased, the accuracy of this procedure falls to chance. This suggests that the activation of each ensembles is associated with some “signature” identifiable in the LFP itself, posing the question of whether an “equivalence relation” exists between ensembles across animals via their LFP signature.

### Statistics of ensemble structure across animals

Finally, we sought to quantify the extent to which the underlying ensemble structure is conserved across animals. We fit LER to data (n=5 mice) over a range of ensemble numbers K, with 10 realizations of LER fitted per K, per mouse. We find that the latent ensembles are disjoint for all animals: for choices of K larger than 3, the cell vectors of distinct ensembles (*σ*) had Pearson correlations near 0 (**Figure S2A**, top). In contrast, we find relatively higher activity correlations between ensembles, as measured by Kendall’s tau (**Figure S2A**, bottom). For K values larger than 3, off-diagonal entries in the cell correlation matrix are consistently weakly to moderately correlated; this correlation increases monotonically and reached τ=0.5 when K=15. This observation validates our choice of K=6 across mice, where activity correlation between ensembles is weakly positive, suggesting some contribution of multiple ensembles to each spike; selecting a larger value for K than the true number of underlying ensembles would artificially split the true ensembles, resulting in spurious ensembles with highly correlated activity. Importantly, we found that irrespective of the number of ensembles specified, the ratio of abGC-dominated ensembles to mGC-dominated ensembles remains relatively fixed at 4:6 across mice (**Figure 4G**). As abGCs themselves represent about 11% of the identified cells in our dataset, and less than 5% of GCs in the DG network (van Praag et al., 2002, Toni et al., 2007, Toni et al., 2008), this result strongly suggests that abGC-dominated ensembles disproportionately drive network activity in the epileptic DG during IEDs. The normalized activity of abGC-dominant vs mGC-dominant ensembles is similar and decreases as K is increased (**Figure S2C**). We also compared the “purity” of the “purest” abGC ensemble to the “purest” mGC ensemble and found that this “maximum purity” (expressed as a fraction from 0 to 1) was increasing with K for both populations, but nearly pure (>90%) mGC ensembles were identified regardless of K, whereas the purest abGC ensemble only contained 80% abGCs at K=20 (**Figure S2D**). This result further suggests that certain mGCs are highly coupled to abGCs and may propagate abGC activity through the network. Finally, we found that the sorted K-dimensional vectors of fractions of abGCs were highly Pearson correlated across mice and across choice of K (**Figure S2E**), suggesting that different animals organize ensembles with similar proportions of abGCs and mGCs. Together, we conclude that different epileptic animals organize ensembles with highly similar statistical properties, suggesting conserved structural and functional motifs underlying the epileptic network.

## Discussion

How macroscale pathological events emerge from microscale changes at the cellular level has been a longstanding open question in the study of TLE (Farrell et al., 2019). Previous imaging and electrophysiological work examining the role of specific cell types in epileptic pathology has shown that inhibitory circuits play a critical role in shaping interictal dynamics in CA1 (Ewell et al., 2015, Muldoon et al., 2015, Miri et al., 2018). However, relatively little is known about how the activity of the DG—the “gating” entry node to the tri-synaptic circuit—shapes downstream excitation-inhibition dynamics *in vivo* through excitatory output during these events. Changes within the DG abGC network have been hypothesized to contribute to the breakdown of the dentate gate which enables seizures to occur (Thind et al., 2008, Krook-Magnuson et al., 2015, Althaus et al., 2019, Danzer, 2018, Cho et al., 2015). This question is highly clinically relevant, as any disease-modifying targeted therapy will depend on a robust understanding of the functional targets of the disease. In this study, we focused on the contribution of abGCs to epileptic DG circuits involved in IEDs and present a novel approach to evaluate the contribution of genetically identified neural populations to IEDs. We recorded abGCs and mGCs during IEDs, which reveal “snapshots” of discrete functional components underlying the epileptic network, and collated these snapshots under a novel generative model framework of ensemble recruitment to deduce the hidden functional organization within these networks.

The results presented here describe how populations of abGC and mGCs within the epileptic DG network differentially participate in macro-level interictal events *in vivo*. Consistent with previous work *in vitro* (Staley and Dudek, 2006, Keller et al., 2010, Feldt Muldoon et al., 2013), we find significant heterogeneity in LFP and calcium responses between IEDs, with distinctly identifiable pure mGC-driven events. The events in which a cell participates constitute a unique fingerprint that is highly informative about the cell’s developmental lineage (abGC vs mGC). We also find significant heterogeneity in the population responses of abGC and mGC cell-types to individual events, as well as in the within-population cell recruitment across cell-type dominated events.

We introduce Latent Ensemble Recruitment (LER), a biologically-motivated generative model of cell recruitment by interictal events. This model represents the first application of Bayesian topic modeling to *in vivo* two-photon calcium imaging data, as well as a novel conceptual framework for relating interictal events to ictogenic circuits: each micro-level population event represents a “snapshot” of a much wider macro-level network, whose functional or synaptic organization is not directly observable, yet whose micro-level calcium response at each time point is. Each one of these snapshots reveals some information about the underlying network, which is hypothesized to be structured on multiple scales to support seizure initiation and propagation (Paz and Huguenard, 2015, de Curtis and Avoli, 2015). While each individual event is relatively uninformative on its own, by accumulating these snapshots in an unsupervised way, it is possible to reconstruct a more complete picture of the underlying macro network structure. When fit to real data, this cell type- and event time-agnostic model discovers micro-ensembles with clear cell-type and temporal organization. This observation suggests that, while the recruitment of any individual cell by IEDs may appear to be random, the recruitment of ensembles exhibits statistical regularities which strongly imply a role for the underlying microcircuits in the generation of epileptiform events. In particular, the regularities which we observe correspond to biological intuitions about the underlying circuits –e.g., the observed repeated activations in time may correspond to a microcircuit entering and remaining in a hyperexcited state for several seconds, such that the circuit is highly likely to be recruited by global events which occur within that window.

Our modeling approach revealed that, abGC-dominated ensembles disproportionately drive network activity in the epileptic DG during IEDs, which is consistent with previous observations. Finally, for small ensemble numbers, the most active ensemble in each IED could be decoded significantly above chance from the power spectrum of the IED alone. This suggests that specific GC ensembles have distinct LFP signatures (see Valero et al., 2017 for another example). While there is no obvious way to equate ensembles across animals, whether these LFP signatures are conserved across animals poses an interesting open question for future work. We hypothesize that there exists an equivalence relation between the LFP spectral signatures of IEDs (whether from the same or different animals); combined with our observations here, this would imply that an equivalence class structure exists for cellular ensembles across animals via their LFP signatures, possibly offering an effective bridge to the micro-macro disconnect. While our results here apply to the adult-born and mature subpopulations of the principal cell population in the DG, our framework of latent ensemble recruitment could motivate future experiments to also examine DG interneurons and mossy cells, which also play critical roles in the epileptic DG circuit, (Hofmann et al., 2016, Bui et al., 2018, Scharfman, 2016) and more generally, to investigate cell type-specific ensemble dynamics in other HPC subregions during inter-ictal and ictal events (Muldoon et al., 2015, Ewell et al., 2015, Miri et al., 2018).

Recent work (Johnston et al., 2020) has reported that rAAVs impair neurogenesis in the adult mouse DG, with significant implications for functional recordings of GCs relying on rAAV for delivery of a genetically encode actuators or sensors. Given the design of our study, we believe these findings have limited relevance to the work presented here: principally, Johnston and colleagues found that cells born two weeks or more prior to viral injection demonstrate no reduction, and in our study induction of Nestin expression by TMX occurred 2-3 weeks prior to injection with AAV1.Syn.GCaMP6f. In addition, the total volume of virus used in this study, 196 nl, is less than 1/5 the volume found to cause death of newborn cells at the same titer. Finally, even at the larger volume of 1000 nl, Johnston et al only observed a “partial effect” of ablation at the viral titer used in this study, 1×10^12^. Most compellingly, during imaging 3 weeks post injection, we empirically see a large and active population of abGCs indelibly labeled within the NestinCreER^T2^ line—if this population has somehow been reduced, this effect would make our observation of an identifiable distinct role for abGCs in TLE even more remarkable.

Identification and characterization of circuit-level functional organization represents a critical step toward a general framework for resolving the micro-macro disconnect in chronic epilepsies (Farrell et al., 2019). Here we provide the first *in vivo* characterization of single cell and microcircuit-level dynamics in multiple cell populations in the epileptic DG and relate them to macroscale events in simultaneously recorded LFP. These results may suggest a new design strategy for closed-loop systems for intervening in epilepsy, based on actively recognizing the patient-specific microcircuit targets for intervention *in situ*. Two-photon calcium imaging is currently the only technique that permits the longitudinal functional characterization of physiological and pathological neural circuits at the cellular level, which is necessary for the dissection of epileptic microcircuits and their recruitment during IEDs *in vivo*. Currently, continuous monitoring of microcircuit dynamics using calcium imaging is not translatable to humans. However, long-term scalp and invasive EEG recording is routinely conducted as part of the pre- and post-surgical evaluation for intractable focal epilepsy syndromes, and is the *de facto* diagnostic and monitoring tool for abnormal brain activity. The input signal to any therapeutic closed-loop intervention system will be electrographic for the foreseeable future, and the critical “inverse problem” for relating electrophysiological observables back to circuit mechanisms is unavoidable. Our work here takes the first steps toward solving this “inverse problem” by connecting signatures in LFP to circuit-level events, which will be essential for the design of new-generation functional closed-loop interventions.

## Acknowledgments

Supported by grants NINDS-1U19NS104590 (A.L., I.S.), NINDS-1R01NS094668 (A.L., I.S.), and the Kavli Foundation (A.L.). We would like to thank P. Maccario for help running IED classification, H. Kim for assisting with vEEG software, and D. Hadjiabadi for comments on the manuscript. We also thank C. Schevon for helpful comments on a previous version of the manuscript.

## Author contributions

F.T.S, Z.L., I.S., and A.L. designed the experiments. F.T.S and Z.L. collected and analyzed the data. F.T.S and W.L. designed electrophysiology hardware components. F.T.S, Z.L., I.S., and A.L. wrote the manuscript.

## Methods

### CONTACT FOR REAGENT AND RESOURCE SHARING

Further information and requests for resources and reagents should be directed to the Lead Contact Attila Losonczy. All unique/stable reagents generated in this study are available from the Lead Contact with a completed Materials Transfer Agreement.

### EXPERIMENTAL MODEL AND SUBJECT DETAILS

#### Animals

All experiments were conducted in accordance with the US National Institutes of Health guidelines and with the approval of the Columbia University Animal Care and Use committee. Male transgenic mice were obtained from The Jackson Laboratory to establish a local breeding colony on a C57BL/6J background: Nestin-CreER^T2^ (JAX:016261) and ROSA26-CAG-stop^flox^-tdTomato Ai9 (JAX:007909). The Nestin-Cre line was crossed with the Ai9 reporter line to express tdTomato in adult-born granule cell populations. Mice were housed in the vivarium on a 12h light/dark cycle, were housed 3-5 mice per cage, and had access food and water *ad libitum*. Mice were housed individually during video-EEG monitoring following kainic acid injection. Mature male and female mice (>8 weeks of age) were used for all experiments.

#### Induction of transgene expression and experimental timeline

All mice were induced at approximately eight weeks of age, two weeks prior to kainic acid (KA) injection. For induction of transgene expression in Nestin-CreER^T2/tdTomato^ mice, 3mg tamoxifen (TMX) (20mg/mL in corn oil/10% ethanol) was injected I.P./day for 5 consecutive days. abGCs were indelibly labelled with tdTomato following injections of TMX that drove expression of Cre in Nestin+ cells. This induction procedure labels approximately 90% of the immature granule cell population expressing doublecortin (Danielson et al., 2016a). Two weeks later, KA was injected into the ventral hippocampus to induce the epilepsy model, and GCaMP6f injected into the dorsal dentate gyrus. Following recovery from injection of KA, mice were placed in video-EEG enabled housing where LFP and behavioral activity were continuous recorded to monitor ictogenesis (Armstrong et al., 2013). Three weeks post-KA injection, mice were habituated to being head fixed under the two-photon microscope, and Ca^2+^ imaging proceeded over the following 1-2 weeks (**Figure 1C**).

#### Surgical procedures

##### Imaging window implant

Recombinant adeno-associated virus carrying the GCaMP6f gene (AAV2/1:hSyn-GCaMP6f) was obtained from Addgene (100837-AAV1) with titer ≥ 1×10^12^. The dorsal dentate gyrus was stereotactically injected using a Nanoject syringe, as described previously (Lovett-Barron et al., 2014, Kaifosh et al., 2013). Injection coordinates were -2.3 mm AP, 1.5 mm ML, and -1.8, -1.65, -1.5 mm DV relative to the cortical surface. 64 nL of virus was injected at each DV location in 32 nL increments at a flow rate of 23 nL/sec. Three days later, mice were then surgically implanted with an imaging window (diameter: 2.0 mm; height: 2.3 mm) over the left dorsal dentate gyrus. Imaging cannulas were constructed by adhering a 2 mm glass coverslip (custom cut by Potomac) to a cylindrical stainless-steel cannula (2 mm diameter x 2.3 mm height) using optical adhesive (Norland). The surgical procedure was very similar to that previously described (Danielson et al., 2016a), the imaging window was implanted 100-200 µm above the hippocampal fissure, providing optical access to the granule cells in the dorsal blade of the DG, and the interneurons and mossy cells in the hilus. Briefly, following induction of anesthesia (Isoflurane: 3.5% induction, 1.5-2.0% maintenance; 1.0 L/min O2) and administration of analgesia (Metacam 5 mg/kg i.p.; Bupivacaine 2 mg/kg s.c.), the scalp was removed, and a 2.0 mm diameter craniotomy centered over the injection location was performed, using a fin-tipped dental drill. The dura was removed, and the underlying cortex aspirated until fibers within the *stratum lacunosum moleculare* were visible. The cannula with window was placed within the aspirated cavity and fixed to the skull with dental cement. A stainless steel straight headpost was then cemented to the skull posterior to the imaging window.

##### Electrode implants

During the window implant surgery, electrodes were implanted to monitor hippocampal local field potentials and neck muscle electromyographic signals. A custom monopolar electrode was constructed from 127 μm Teflon coated stainless-steel wire (A-M Systems), and inserted near the location of the KA injection in the ventral hippocampus (AP: -3.28 mm, ML: 2.75 mm, DV: -2.5 mm) ipsilateral to the imaging window to monitor and record hippocampal local field potentials. The location of the depth electrode was chosen based on previous studies showing that spontaneous seizures in the KA epilepsy model typically arise from the hippocampal formation ipsilateral to the KA injection site (Bragin et al., 1999b). A stainless-steel jewelers screw was placed in the contralateral frontal bone for a ground electrode. To record electromyographic signals, a second stainless-steel screw was inserted in the occipital bone through overlying cervical trapezius muscle. Electrode wires were connected to a custom plug (Mill-Max strip connector) that was then cemented to the headpost. Following recovery from the implant procedure, mice underwent 24 h video-EEG monitoring for seizure detection.

##### Kainic acid injection

KA (AG Scientific, USA) was dissolved in sterile phosphate buffered saline with a final concentration of 20 mM. Using procedures described above, mice were stereotaxically injected while under isoflurane anesthesia, with 50 nL kainic acid unilaterally into a single location within the ventral hippocampus (AP: -3.28 mm, ML: 2.75 mm, DV: -3.4 mm). The location of the KA injection was chosen to allow for chronic 2-photon imaging of the dorsal dentate gyrus ipsilateral to the KA injection. Intra-ventral hippocampal KA injection results in epileptogenesis and seizure profiles similar to that found in the dorsal hippocampal KA model (Zeidler et al., 2018).

#### In vivo 2-photon imaging of dentate gyrus

We used the same imaging system as described previously (Zaremba et al., 2017, Danielson et al., 2016b). All imaging was conducted using a two-photon microscope equipped with an 8 kHz resonant scanner (Bruker). Approximately 50-100 mW of laser power under the objective was used for excitation (Ti:Sapphire laser, (Chameleon Ultra II, Coherent) tuned to 920 nm), with adjustments in power levels to accommodate varying window clarity. To optimize light transmission, we adjusted the angle of the mouse’s head using two goniometers (Edmund Optics, +/-10-degree range) such that the imaging window was parallel to the objective. A piezoelectric crystal was coupled to the objective (Nikon 40X NIR water-immersion, 0.8 NA, 3.5mm WD), allowing for rapid displacement of the imaging plane in the z-dimension. We continuously acquired red (tdTomato) and green (GCaMP6f) channels separated by an emission cube set (green, HQ525/70m-2p; red, HQ607/45m-2p; 575dcxr, Chroma Technology) at 512 x 512 pixels covering 225 μm x 225 μm, at 8-30 Hz (dependent on number of planes imaged) with photomultiplier tubes (green GCaMP fluorescence, GaAsP PMT, Hamamatsu Model 7422P-40; red tdTomato fluorescence, multi-alkali PMT, Hamamatsu R3896). A custom dual stage preamp (1.4 x 10^5^ dB, Bruker) was used to amplify signals prior to digitization.

For all experiments, mice were head-fixed and ran freely on a 2m long treadmill belt. We habituated the mice to the head-fixed condition for at least one hour per day over three days prior to the beginning of the experiment.

#### Electrophysiology recording

All mice were implanted with a hippocampal LFP electrode and imaging window. During two-photon imaging, hippocampal LFP was recorded so that electrographic events could be correlated and analyzed with calcium imaging data. Electrophysiology signals were acquired with a multichannel digital recording system (Intan Technologies, USA) at 25kHz and synchronized with the frame-start signal of the microscope. While not being imaged, mice were routinely monitored for interictal and seizure events using a custom continuous video-EEG system previously described (Armstrong et al., 2013, Krook-Magnuson et al., 2013), though we used an analog multichannel recording system customized to record up to 16 mice simultaneously (NeuraLynx, USA). Briefly, the hippocampal LFP signal and video were acquired by a PC running a custom MatLab seizure recording and detection algorithm. EEG signals were acquired for offline determination of IED, and seizure frequency and duration.

#### Histology

Mice were deeply anesthetized with isoflurane (5% in O2; 1L/min) and perfused with 0.1M phosphate buffered saline (PBS) followed by 4% paraformaldehyde (PFA) in PBS. Brains were dissected and post-fixed in 4% PFA overnight. 40µm sections were cut throughout the hippocampus on either a cryostat or freezing sliding microtome. Free-floating sections were first stained with a blue fluorescent Nissl stain (NeuroTrace, ThermoFisher, USA) and then mounted on slides for imaging. Widefield images for gross lesion quantification were acquired with a Nikon AZ100 Multizoom epifluorescence slide scanning microscope, while high resolution images were acquired with a Nikon A1R laser scanning confocal microscope.

### QUANTIFICATION AND STATISTICAL ANALYSIS

#### Ca^2+^ imaging data preprocessing

All imaging data were pre-processed using the SIMA software package (Kaifosh et al., 2014). Motion correction was performed using whole frame registration. In cases where motion artifacts were not adequately corrected, the affected data were discarded from further analysis. We used the Suite2p software package (Pachitariu, 2017) to identify spatial masks corresponding to neural regions of interest (ROIs) and extracted associated fluorescence signal within these spatial footprints, correcting for cross-ROI and neuropil contamination. Identified ROIs were curated post-hoc using the Suite2p graphical interface to exclude non-somatic components. ROI detection with Suite2p is inherently activity-dependent, and so for each session, we detected only a subset of neurons that were physically present in the FOV. To track ROIs across imaging sessions, all recordings of the same FOV were first concatenated before calculating spatial masks with Suite2p.

#### Running modulation calculation

For each calcium trace triggered on an event, the mean of the pre-event activity was subtracted from the mean of the post-event activity (mean ΔF/Fpost-event – mean ΔF/Fpre-event) to calculate the response magnitude. To calculate whether a cell reliably changes its activity on an event type, the response magnitudes for that event type were sampled with replacement, and the mean response magnitude was calculated. We repeated this calculation for 3000 bootstrap resampling iterations, to construct a 99% confidence interval (CI). The cell was determined to have be significantly negative response if the CI was less than 0, significantly positive response if the CI was greater than 0, and non-significant response if the CI contained 0.

#### Classification of cell type and epileptiform events

IEDs were determined from single-channel LFP using a semi-supervised approach. First, the LFP signal was aligned with each imaging frame using an external frame-trigger ADC channel. The magnitude spectrum of the LFP snippet corresponding to each imaging frame was calculated using the Fast Fourier Transform (FFT). The magnitude spectrum was chosen so that IEDs could be detected regardless of phase within the frame. We then trained an Online Kernelized Perceptron (using a Gaussian radial basis function kernel with σ = 1) to recognize frames containing an IED by hand-annotating a small number of LFP snippets containing IEDs, allowing the classifier to classify all the snippets in the session based on the hand-labeled examples, and updating annotations based on the classifier’s feedback. This classification procedure was conducted according to an iterative feedback process, where human input would train the algorithm while correcting the algorithm’s mistakes, so that each session is hand-curated, but the amount of hand curation necessary is minimized. We verified the internal consistency of this algorithm quantitatively by plotting the 1/f corrected average IED power spectrum (Fig 2B); we find the classified IEDs share common spectral characteristics. An online perceptron-type algorithm was chosen because it permits efficient human-supervised training with immediate feedback after each labeled example and highly accurate phase-agnostic classification with a small number of labeled samples. A kernelized algorithm was chosen so that nonlinear relationships between IEDs and the magnitude spectrum could be captured.

#### Latent Ensemble Recruitment Modelling

Synchronous network events were identified using spectral analysis of a simultaneously collected single-channel LFP recording in ipsilateral CA1. The “responsiveness” Δ of a cell to an event is defined as the cell’s mean activity in a fixed window (e.g., 3 s) pre-event subtracted from the cell’s mean activity in the same window post-event.

Existing data from *in vitro* physiological studies (Muldoon et al., 2015) have suggested that epileptiform activity is associated with spatially clustered, temporally coactivated neural ensembles. Our model (**Figure 4A**,**B**) formalizes a weaker version of this hypothesis: that each interictal spike recruits a sparse set of pre-existing ensembles within the network, and each ensemble activates a subset of its constituent cells. Specifically, we assume the following generative process for a dataset consisting of *M* events in a population of *N* cells organized into *K* ensembles: Once for the entire network, we sample the ensembles *σ*_*k*_ for each *k* from a Dirichlet distribution parameterized by sparsity-controlling hyperparameter *β*. This gives us *K* ensembles, where each ensemble *σ*_*k*_ is a probability vector over the *N* cells in the network, whose entries correspond to the strength of each cell’s association with that ensemble. For each event *i* = 1,…, *M*, we sample ϑ*i*, a sparse probability vector over ensembles, from another Dirichlet distribution parameterized by hyperparameter *α*, representing the degree to which each ensemble is recruited by that event. Then, for each cell, we sample the indicator variable *z*_*ij*_ from a Bernoulli distribution parameterized by *p* = ∑_*k*_ *θ*_*ik*_ *σ*_*kj*_, which represents whether or not that cell was recruited to that event. We simultaneously sample an ensemble ζ from the multinomial Multinomial(±φ), which gives the ensemble from which the cell *j* was recruited. Finally, we draw our observed responsiveness Δ from a Gaussian, either 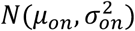 *or* 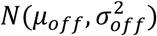 depending on the value of *z*_*ij*_.

