## Supplemental information for "Adult-born granule cells support pathological microcircuits in the chronically epileptic dentate gyrus"

**Figure S1**

**A.** Average *z*-scored ΔF/F response to locomotion onset, offset, and IED , where 'responsiveness' is defined as mean ΔF/F_post – mean ΔF/F_pre, where 'pre' and 'post' refer to 3 second windows before and after the event onset, by imaged genetically identified cell types (subjects n = 5 mice for mGCs (n = 2162), abGC (n = 277). Each dot represents the average population response of one mouse. We find no significant difference in running start ($p$ = 0.39) or running stop ($p$ = 0.08) responsiveness between the abGC and mGC populations, but we do find that the abGC population is more responsive to IEDs by a small but significant margin compared to the mGC population ($p < 1\times10^{-4}$, Mann-Whitney *U*-test).

**B.** PSTHs of single-cell responses to "most informative" IEDs from 10 minute-recording from one mouse. Single-cell responses to pro-mGC (left) and pro-mGC (right) events exhibit high levels of heterogeneity despite population trends.

**C.** Population responses to pro-mGC and anti-mGC events identified in *Figure 3*. A two-way analysis of variance yielded a main effect for IED class (pro-mGC vs anti-mGC), $F(3,54) = 14.11$, $p=6\times10^{-7}$, such that pro-mGC IEDs significantly positively modulated mGCs compared to anti-mGC IEDs. The main effect of population was non-significant, $F(3,54)=0.13$, $p=0.71$, which suggests that our observations cannot be explained by intrinsic differences in responsiveness between the two populations independent of the type of IED. However, the interaction effect was significant, $F(3,54) = 10.29$, $p=0.002$, indicating a crossing over effect, i.e., that pro-mGC IEDs significantly positively modulated mGCs over abGCs.

**Figure S2. Ensemble statistics are conserved across animals.** We sought to quantify the extent to which the ensemble structure learned for one mouse generalizes across animals. We fit LER to data from N=5 epileptic mice with fields of view in the GCL over a range of ensemble numbers K, with 10 realizations of LER fitted per K per mouse.

A. Relationship between LER and LDA. Top: Analogy between Latent Dirichlet Allocation (LDA) for topics and LER for ensembles. LDA finds topics interpreted as "sports-related" or "business-related." LER finds ensembles of co-occurring neuronal activations that can be interpreted as "abGC-related" or "mGC-related". Bottom: Base LER graphical model (left) and reduction to LDA (right) via point estimate of z (hatched).

**B.** Average off-diagonal correlations. The average off-diagonal correlation measures the degree to which two distinct ensembles, chosen at random, are similar. Top: correlations by cell, measured by Pearson's ρ. Except when K is small (<4), off-diagonal entries in the cell correlation matrix are very close to 0, supporting the previous observation that the latent ensembles underlying the network are disjoint. Bottom: correlations by activity, measured by Kendall's τ. Except when K is small, off-diagonal entries in the cell correlation matrix are weakly positive, suggesting a "primary" responding ensemble to each event with some contribution by several others; this correlation increases monotonically and reached $\tau = 0.5$ when K=15. We choose $\tau = 0.5$ as the cutoff correlation for non-spurious ensembles; any number of ensembles smaller than 15 results in similar qualitative observations. We encode our prior biological knowledge and intuitions about the problem in a novel graphical model (LER, left), and subsequently show that this biological descriptive model can be transformed into a model which we know how to perform inference on (LDA, right).

C. Mean normalized activity of abGC- vs mGC-dominated ensembles, plotted as a function of K.

D. The "purity" of the "purest" abGC ensemble to the "purest" mGC ensemble, as a function of K. This "maximum purity" (expressed as a fraction from 0 to 1) is increasing with K for both populations, but nearly pure (>90%) mGC ensembles can be identified regardless of K, whereas the purest abGC ensemble only contains 80% abGCs at K=20, suggesting that certain mGCs are highly coupled to abGCs and may propagate abGC activity through the network.

E. Correlation matrices of the sorted K-dimensional composition vectors (where each entry is the % of abGCs in an ensemble, sorted from fewest to most). These vectors are highly Pearson correlated across mice and across choice of K, suggesting that different animals organize ensembles with similar proportions of abGCs and mGCs.

**Table S1**

**Per subject Suite2p extracted ROIs**

| | **mGC** | **abGC** | **Total** | **% abGC** |
|---|---|---|---|---|
| FS201 | 142 | 39 | 181 | 21.5 |
| FS202 | 218 | 48 | 266 | 18.0 |
| FS203 | 428 | 39 | 467 | 8.4 |
| FS204 | 763 | 41 | 804 | 5.1 |
| FS205 | 613 | 111 | 724 | 15.3 |
| **Total** | 2164 | 278 | 2442 | |
| **Mean** | 432.8 | 55.6 | 429.5 | 13.3 |
| **SD** | 260.9 | 31.2 | 273.6 | 6.8 |
| **N** | 5 | 5 | 5 | 5 |
| **SEM** | 116. 7 | 13.9 | 122.4 | 3.1 |

**Figure S1**

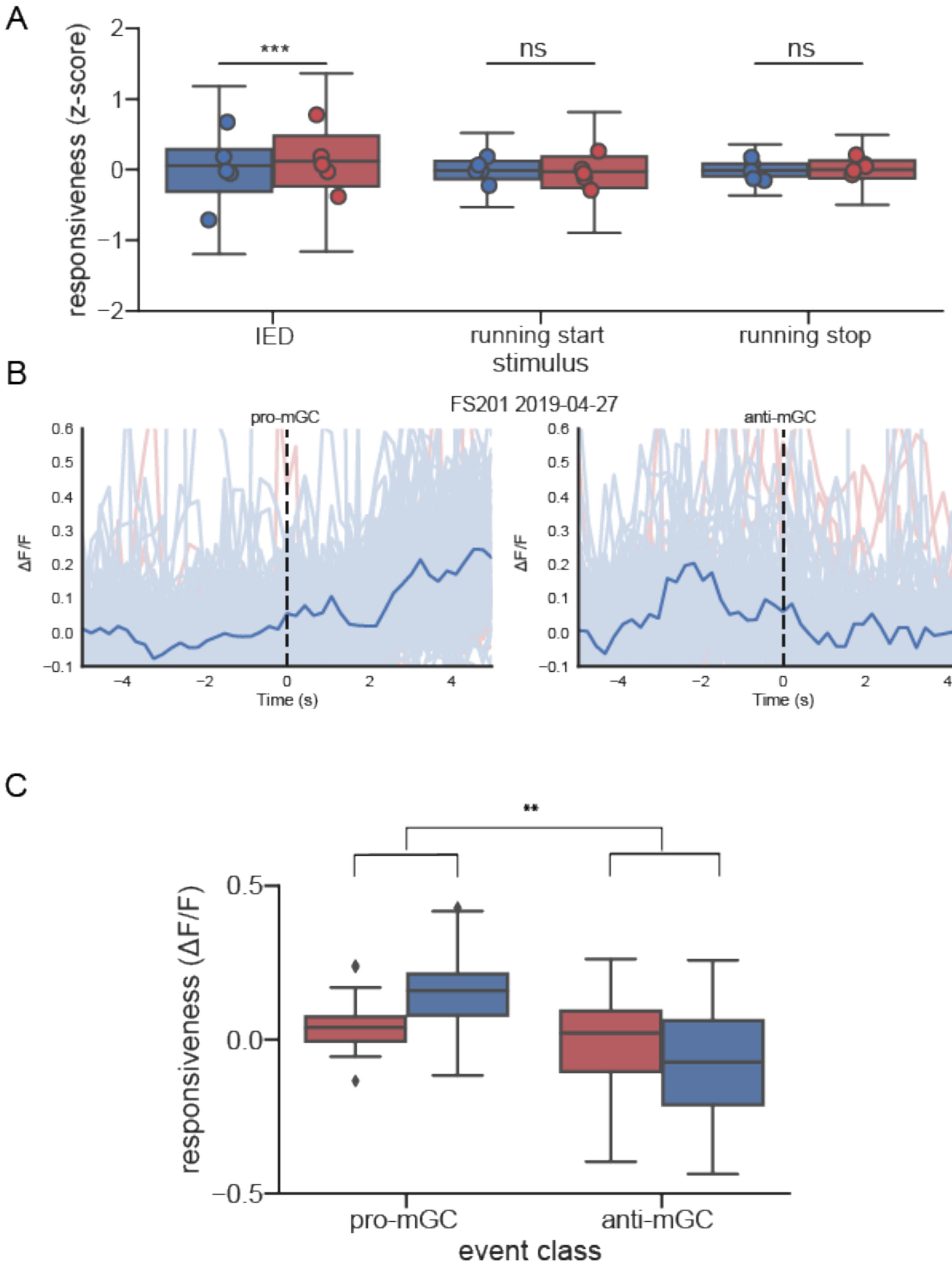
A
***
ns
ns
responsiveness (z-score)
2
1
0
−1
−2
IED
running start
running stop
stimulus
B
FS201 2019-04-27
pro-mGC
anti-mGC
ΔF/F
0.6
0.5
0.4
0.3
0.2
0.1
0.0
−0.1
−4
−2
0
2
4
Time (s)
C
**
responsiveness (ΔF/F)
0.5
0.0
−0.5
pro-mGC
anti-mGC
event class

**Figure S2**

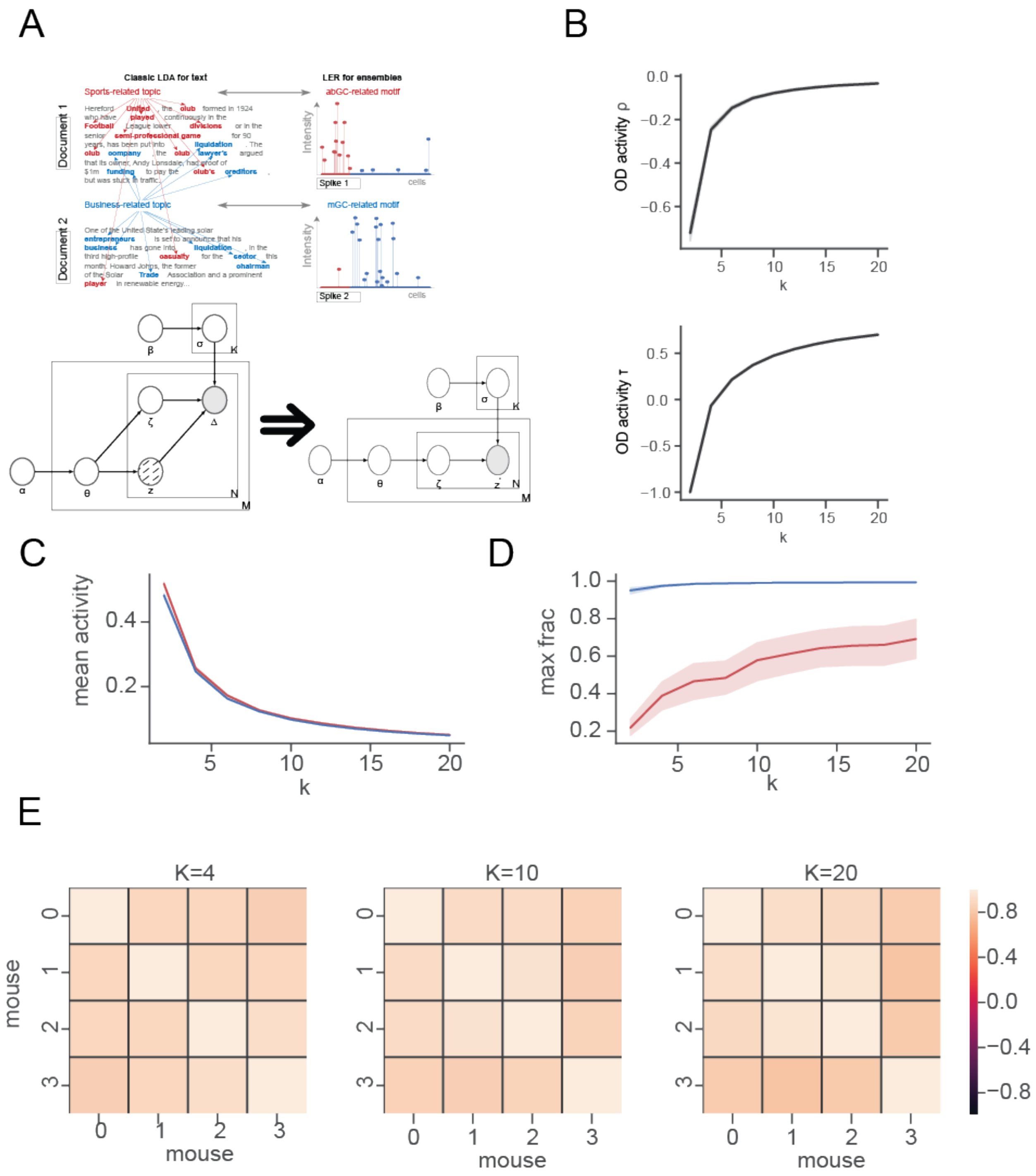
A
B
C
D
E
Classic LDA for text
LER for ensembles
Sports-related topic
abGC-related motif
Business-related topic
mGC-related motif
Document 1
Document 2
Intensity
cells
Spike 1
Spike 2
OD activity ρ
OD activity τ
k
mean activity
max frac
K=4
K=10
K=20
mouse
